# Persistence of an infectious form of SARS-CoV-2 post protease inhibitor treatment of permissive cells in vitro

**DOI:** 10.1101/2023.12.20.572655

**Authors:** Manoj S. Nair, Maria I. Luck, Yaoxing Huang, Yosef Sabo, David D. Ho

## Abstract

Reports have described SARS-CoV-2 rebound in COVID-19 patients treated with nirmatrelvir, a 3CL protease inhibitor. The cause remains a mystery, although drug resistance, re-infection, and lack of adequate immune responses have been excluded. We now present virologic findings that provide a clue to the cause of viral rebound, which occurs in ∼20% of the treated cases. The persistence of an intermediary form of infectious SARS-CoV-2 was experimentally documented in vitro after treatment with nirmatrelvir or another 3CL protease inhibitor, but not with a polymerase inhibitor, remdesivir. This infectious intermediate decayed slowly with a half-life of ∼1 day, suggesting that its persistence could outlive the treatment course to re-ignited SARS-CoV-2 infection as the drug is eliminated. Additional studies are needed to define the nature of this viral intermediate, but our findings point to a particular direction for future investigation and offer a specific treatment recommendation that should be tested clinically.

---

Paxlovid, an FDA approved drug to treat symptomatic SARS-CoV-2 infection in elderly and high-risk individuals^1^, is an oral combination of nirmatrelvir, an inhibitor of the main protease (3CL) of SARS-CoV-2, and ritonavir, a CY3PA inhibitor that boosts the plasma concentration of nirmatrelvir^2^. Another protease inhibitor approved for clinical use in Japan is ensitrelvir (also known as S-217622)^3^.

A number of COVID-19 patients receiving the recommended 5-day course of Paxlovid (300mg nirmatrelvir/100mg ritonavir every 12h) became symptomatically better and virus negative only to have a rebound of detectable SARS-CoV-2 again 2-8 days later^4^. Many of these cases also had a recurrence of symptoms, albeit mild. The Centers for Disease Control and Prevention issued a health advisory because of concerns for forward transmission of the virus during its recrudescence^5^, as another case series was reported^6^. Several retrospective studies suggested the prevalence of “Paxlovid rebound” was low at ∼1-2%^7-9^, but a recent, comprehensive, and prospective study of 72 patients treated with Paxlovid found that 20.8% (n=15) developed sustained viral rebound^10^, as many medical practitioners have noted anecdotally. This report also showed the low prevalence found in prior retrospective studies was largely due to inadequate post-treatment sampling to detect the virus. In addition, some studies suggested viral relapse was common in untreated patients^11,12^, but these were descriptions of viral “blips” that were not sustained. Moreover, the infrequency (0.7%) of SARS-CoV-2 rebound in the absence of therapy is well documented in a large study (n=999) of infected individuals who were closely monitored^13^. An explanation for the viral relapse remained elusive^14^, although viral resistance to nirmatrelvir, inadequate adaptive immunity, and re-infection were ruled out as possible explanations^4,6,10,15,16^. In this study, we probed the underlying cause of the viral rebound by assessing the persistence of infectious SARS-CoV-2 in several permissive cell lines after treatment with high doses of nirmatrelvir or ensitrelvir in vitro.

To ensure maximum inhibition of SARS-CoV-2, we first determined the inhibitory potency of the two protease inhibitors (nirmatrelvir, ensitrelvir) and a polymerase inhibitor (remdesivir, as control) in three different permissive cell lines – Huh7-ACE2, A549-ACE2 and Vero-TMPRSS2. All the drugs showed robust virus inhibition in all the cell lines (**Figure S1**), as indicated by their 50% and 99% inhibitory concentrations (IC_50_ and IC_99_) (**Figure S1**). These results enabled the follow-up in vitro experiments to assess serially the persistence of replication-competent forms of SARS-CoV-2 post exposure to drug concentrations ≥10-fold higher than the IC_99_.

In the pilot experiment, we examined the persistence of infectious virus in Huh7-ACE2 for three consecutive days after treatment with each drug (nirmatrelvir, ensitrelvir, or remdesivir) at 10x IC_99_ 6h prior to inoculation with SARS-CoV-2 /USA/WA1 bearing the mNeonGreen reporter gene (SCoV-2/mNG)^17^ at 0.5 multiplicity of infection (MOI). At 24h, 48h, and 72h post infection, batches of cells were washed, counted, and then subjected to a serial 3-fold titration for infectious virus starting with 12,500 cells per well into Vero-TMPRSS2 indicator cells free of drug. End-point titers of infectious forms of SARS-CoV-2 showed an expected time-dependent decline post nirmatrelvir or ensitrelvir treatment (**Figure 1A**). However, infectious virus was detectable at all time points, at least in some of the wells. Using linear regression analysis of data, the decay half-lives of the infectivity were calculated to be 23.9h for nirmatrelvir and 26.7h for ensitrelvir. In distinct contrast, remdesivir-treated cells had no measurable infectivity at all time points assessed (**Figure 1A**). This initial finding suggested that while nirmatrelvir or ensitrelvir could block the main viral protease^2,18^, a replication-competent form of the virus can persist intracellularly long enough to re-initiate infection once the drug is removed.

**Figure 1.**
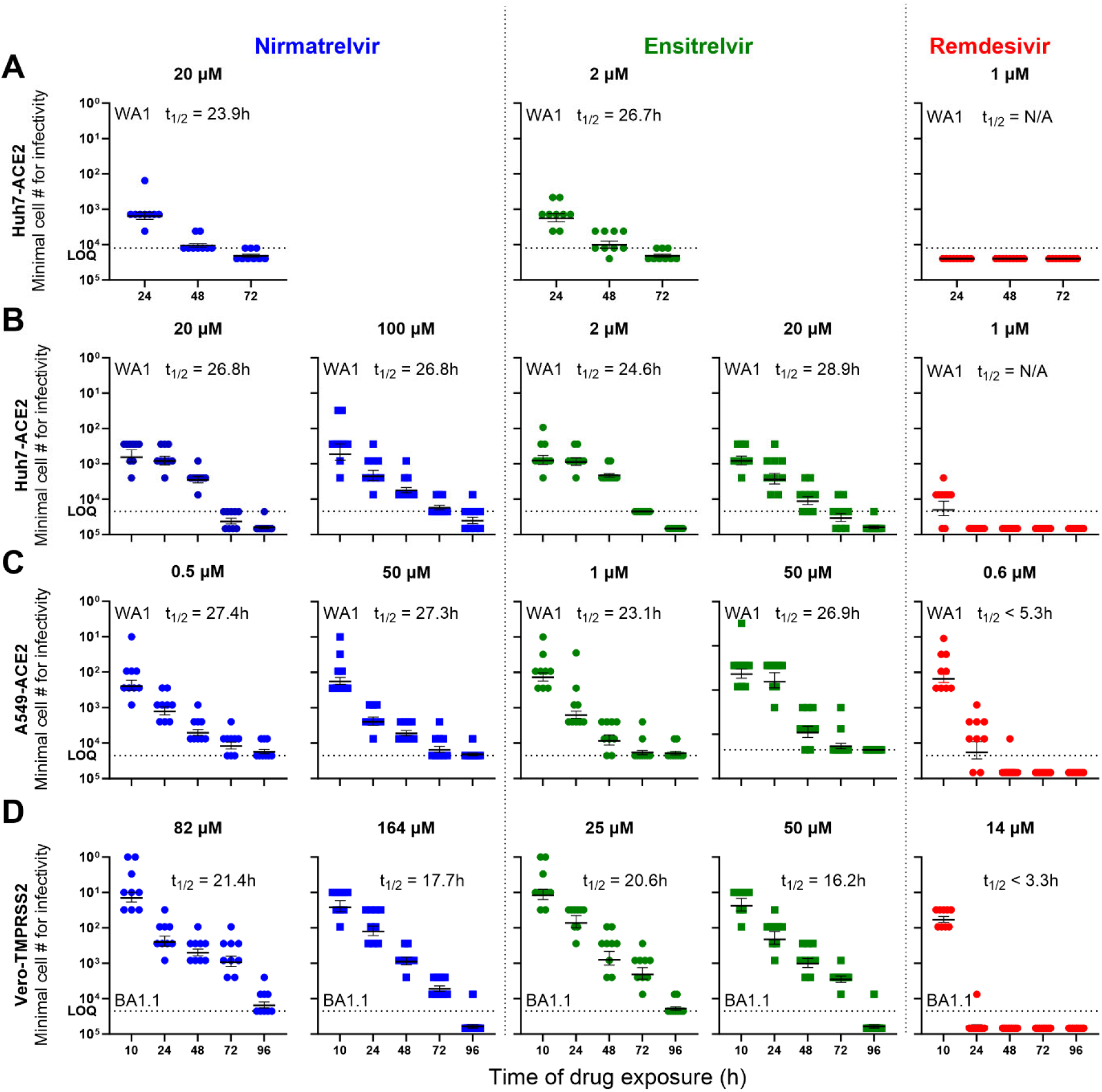
SARS-CoV-2 persistence in cells pre-treated or concurrently treated with protease or polymerase inhibitor at doses 10x IC_99_ or higher. Minimum number of cells retaining infectious virus present post removal of drug treatment at indicated time points are shown in each panel. (**A**) Infectivity decay in Huh7-ACE2 when treated 6h prior to infection, (**B**) Infectivity decay in Huh7-ACE2 when treated concurrently (within 10 min of infection), (**C**) Infectivity decay in A549-ACE2 cells with concurrent treatment, and (**D**) Infectivity decay of Omicron BA.1.1 in Vero-TMPRSS2 cells with concurrent treatment. The dotted line indicates the upper limit of input cells (maximum cell number from which endpoint titration was performed at 3-fold dilutions) for the assay. Inset text shows the drug names and half-life of the decay of infectious form of the virus after treatment with the drug at specified concentrations. Results for nirmatrelvir are shown in blue, ensitrelvir in green, and remdesivir in red.**See also Figure S1 and Figure S2**.

Next, we repeated a similar experiment but with more timepoints assessed as well as higher concentrations of protease inhibitors applied. Huh7-ACE2 cells were again infected with SCoV-2/mNG at 0.5 MOI and concurrently treated with nirmatrelvir, ensitrelvir, or remdesivir at concentrations of 10x IC_99_ or higher. As above, batches of cells were serially washed and harvested to measure the number of cells required to yield infectious virus. Cells treated with either nirmatrelvir or ensitrelvir retained an infectious form of SARS-CoV-2 at the first three time points (10h, 24h and 48h) (**Figure 1B**). By 72h, only 10-30% of replicates yielded detectable infectious virus from cells treated with either of the protease inhibitors. Even at 96h post nirmatrelvir treatment, a minor proportion of replicates (1/9) yielded infectious virus. Again, the rates of infectivity decay were calculated to be 26.8h and 24.6-to-28.9h for nirmatrelvir and ensitrelvir, respectively **(Figure 1B**). Once again, the infectivity decay in cells treated with remdesivir was substantially faster, with no infectious virus detected by 24h.

To exclude cell-type-specific effects, we performed a similar experiment using the same virus (SCoV-2/mNG) but a different cell line, human lung carcinoma-derived alveolar epithelial cells (A549-ACE2). Both nirmatrelvir and ensitrelvir were more potent in A549-ACE2 cells than in Huh7-ACE2 cells (**Figure S1**); however, the infectivity decay results obtained in A549-ACE2 cells (**Figure 1C**) were comparable to those in Huh7-ACE2 cells. The calculated infectivity decay half-lives were 27.3h to 27.4h for nirmatrelvir, 23.1h to 26.9h for ensitrelvir, and <5.3h for remdesivir.

“Paxlovid rebound” was first reported^4^ when the Omicron variant of SARS-CoV-2 was most prevalent. We therefore conducted another similar experiment using Omicron BA.1.1 in yet another cell line, Vero-TMPRSS2. Cells were concurrently inoculated with the virus (0.5 MOI) and treated with 10x IC_99_ or higher concentrations of nirmatrelvir, ensitrelvir, or remdesivir. Similar patterns of infectivity decay were observed once more (**Figure 1D**). Calculated decay rates were slightly faster in this experiment, with half-lives of 17.7h to 21.4h for nirmatrelvir and 16.2h to 20.6h for ensitrelvir. The infectivity decay for remdesivir was again substantially faster, with half-life of <3.3h and no detectable infectious virus after 24h of treatment.

We then tested a third protease inhibitor, GC-376, that was reported to inhibit SARS-CoV-2 in vitro^19^. Dose titration on this investigational compound was again assessed (**Figure S2**) in Huh7-ACE2 and Vero-TMPRSS2 cells. Both cell lines were inoculated with the virus and treated with nirmatrelvir or ensitrelvir at concentrations of ∼10x IC_99_, as previously described. The infectivity decay in both cell lines resembled those of nirmatrelvir and ensitrelvir shown in **Figure 1**, with calculated half-lives of 28.1h for WA1 strain in Huh7-ACE2 cells and 20.3h for the Omicron BA.1.1 strain in Vero-TMPRSS cells (**Figure S2**). Therefore, treatment with all three protease inhibitors in vitro led to similar persistence of an infectious form of SARS-CoV-2 that decays slowly with a half-life of approximately 1 day for the WA1 strain and slightly shorter for Omicron BA.1 or BA.1.1. In contrast, treatment with a polymerase inhibitor resulted in a rapid loss of infectious SARS-CoV-2 in vitro.

We further probed this persistence phenomenon by examining levels of SARS-CoV-2 genomic RNA (gRNA) and nucleocapsid protein (NP) in infected cells treated with nirmatrelvir or remdesivir. Since nirmatrelvir treatment in the clinical setting starts post infection, we modified the in vitro experiment to mimic this scenario. Huh7-ACE2 cells were infected with SCoV-2/mNG at 0.5 MOI for 6 h before the cells were washed and supplemented with nirmatrelvir or remdesivir at concentrations of 10x IC_99_. At 24h, 48h, and 72h post, samples of cells were washed and subjected to endpoint infectivity titration, as well as fixed for simultaneous imaging of viral gRNA by single-molecule RNA FISH (smFISH)^20^ and NP by immunofluorescence.

Cells harboring infectious virions were again detected at all time points from 24h to 72h post nirmatrelvir treatment (**Figure 2A**). Not surprisingly, the observed infectious titers were higher than when an equivalent dose of nirmatrelvir was administered within 10 min of virus inoculation (**Figure 1B**), but the infectivity decay half-lives were similar (24.6h versus 26.8h). The number of cells harboring infectious virus after remdesivir treatment was significantly lower at all time points (**Figure 2A**) by about two orders of magnitude. In the imaging studies, viral gRNA and NP were visibly more abundant at all time points in cells treated with nirmatrelvir than those treated with remdesivir (**Figure 2B** and **Figure S3**). This subjective observation was confirmed when the fluorescence signal intensities were quantified. Viral gRNA levels were indeed significantly higher at 48h and 72h post nirmatrelvir treatment compared to levels post remdesivir treatment (**Figure 2C**). Likewise, significantly higher levels of NP were detected in nirmatrelvir-treated cells at 48h and 72h (**Figure 2D**). While these imaging results seemingly tracked nicely with the infectivity decay data, their causal relationship is unknown.

**Figure 2.**
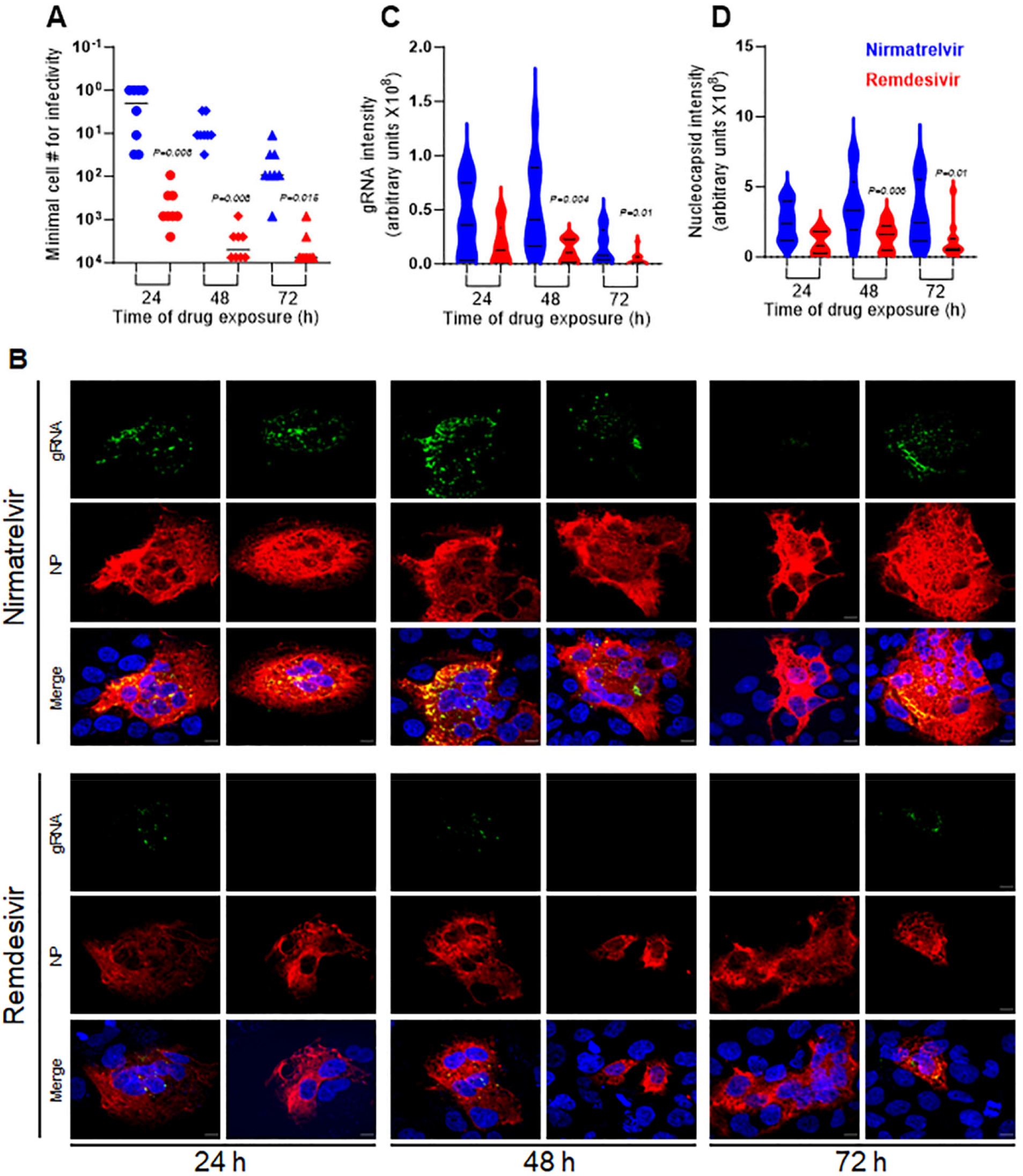
Persistence of infectious SARS-CoV-2 as well as gRNA and NP in Huh7-ACE2 cells after drug treatment. Huh7-ACE2 were infected with SARS-CoV-2 at 0.5 MOI for 6h after which the virus was removed and replaced with growth media supplemented with either 20 μM nirmatrelvir or 1 μM remdesivir. (**A**) Infectivity decay post removal of nirmatrelvir or remdesivir. (**B**) Representative images to demonstrate the persistence of gRNA (green) and viral NP (red) at 24h, 48h and 72h post nirmatrelvir or remdesivir treatment. Cell nuclei were stained with DAPI. Scale bar = 10 μm. Violin plots of the signal intensities for intracellular viral gRNA (**C**) and NP (**D**) in infected cells (collected from **Figure S3**). The exact p values for significant diferences observed between nirmatrelvir and remdesivir are shown.**See also Figure S3**.

In conclusion, our studies on SARS-CoV-2-infected cells in vitro suggest that there is an intermediary form of the virus that is blocked at the stage of polypeptide cleavage by nirmatrelvir or ensitrelvir. The nature of this viral intermediate is yet unclear, but it decays slowly with a half-life of approximately one day. When its persistence outlives the 5 days of treatment, SARS-CoV-2 infection could be re-ignited as the drug is eliminated. This virologic hypothesis alone could explain “Paxlovid rebound” without implicating drug resistance, re-infection, or the immune system. This proposed explanation is also consistent with the observation that the viral rebound is more frequent in patients who were treated early^10^ when the viral load is highest^13^. However, additional molecular and cellular studies are clearly needed to define, more precisely, the nature of the viral intermediate blocked at polypeptide cleavage by inhibitors of 3CL protease. Nevertheless, if our in vitro findings are reflective of the in vivo situation in patients, extending the course of nirmatrelvir treatment by 3-to-5 days should lower the probability of a viral rebound by 8-to-32-fold. This treatment recommendation should be evaluated by clinical studies.

## Acknowledgements

This study was supported by funding from Andrew & Peggy Cherng, Samuel Yin, Barbara Picower and the JPB Foundation, Roger & David Wu, and the Bill and Melinda Gates Foundation. D.D.H., and Y.S are supported by funding from NIAID (1U19AI1711401).

## Author contributions

D.D.H., Y.H., and Y.S. conceptualized the study, M.S.N. and Y.H. performed the virological and dose-response experiments, M.I.L. and Y.S. performed the smRNA-FISH and microscopy experiments. M.S.N., Y.H., Y.S. and D.D.H. wrote the manuscript.

