## Supplementary Information for "Persistence of an infectious form of SARS-CoV-2 post protease inhibitor treatment of permissive cells in vitro"

#### STAR★METHODS

Detailed methods are provided in the online version of this paper and include the following:

- **KEY RESOURCES TABLE**
- **RESOURCE AVAILABILITY**
    - Lead Contact
    - Materials Availability
    - Data and Code Availability
- **EXPERIMENTAL MODEL DETAILS**
    - Cell lines and virus strains
- **EXPERIMENTAL PROCEDURE DETAILS**
    - Dose-response inhibition of virus
    - Determination of minimal cell number for recovering infectivity
    - Single molecule RNA FISH and immunofluorescence
    - Microscopy
    - Viral genomic RNA and nucleocapsid quantification
- **QUANTITATION AND STATISTICAL ANALYSIS**
    - Linear regression analysis to calculate half-life of the infectious form
    - Statistical analysis for determination of significance of dataset

### Supplementary Information

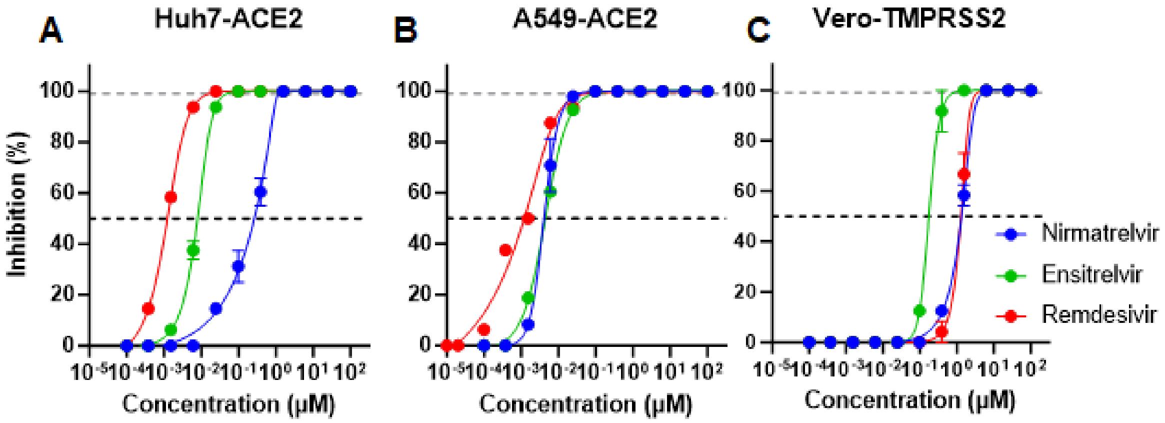


D

| Virus | Target Cell | Nirmatrelvir | | Ensitrelvir | | Remdesivir | |
|---|---|---|---|---|---|---|---|
| | | Inhibitory conc (µM) | | Inhibitory conc (µM) | | Inhibitory conc (µM) | |
| | | $IC_{50}$ | **$IC_{99}$** | $IC_{50}$ | **$IC_{99}$** | $IC_{50}$ | **$IC_{99}$** |
| ic-SARS-CoV-2 /WA1-mNG) | Huh7-ACE2 | 0.266 | **1.789** | 0.008 | **0.186** | 0.001 | **0.073** |
| | A549-ACE2 | 0.004 | **0.035** | 0.004 | **0.090** | 0.001 | **0.060** |
| Omicron BA1.1 | Vero-TMPRSS2 | 1.346 | **8.203** | 0.176 | **2.533** | 1.479 | **6.705** |

**Figure S1: Dose-response inhibition of SARS-CoV-2 by protease and polymerase inhibitors.** Dose titration of drugs in (A) Huh7-ACE2 cells infected with WA1 (ic-SARS-CoV-2/WA1-mNG), (B) A549-ACE2 cells infected with WA1 (ic-SARS-CoV-2/WA1-mNG) and (C) Vero-TMPRSS2 cells infected with Omicron BA.1.1 virus. All viruses were used at MOI 0.5 to obtain the $IC_{50}$ and $IC_{99}$ of inhibition of the proteases and polymerase inhibitor. (D) Table showing potency of each drug in panel A-C in respective cell lines. **See also Figure 1.**

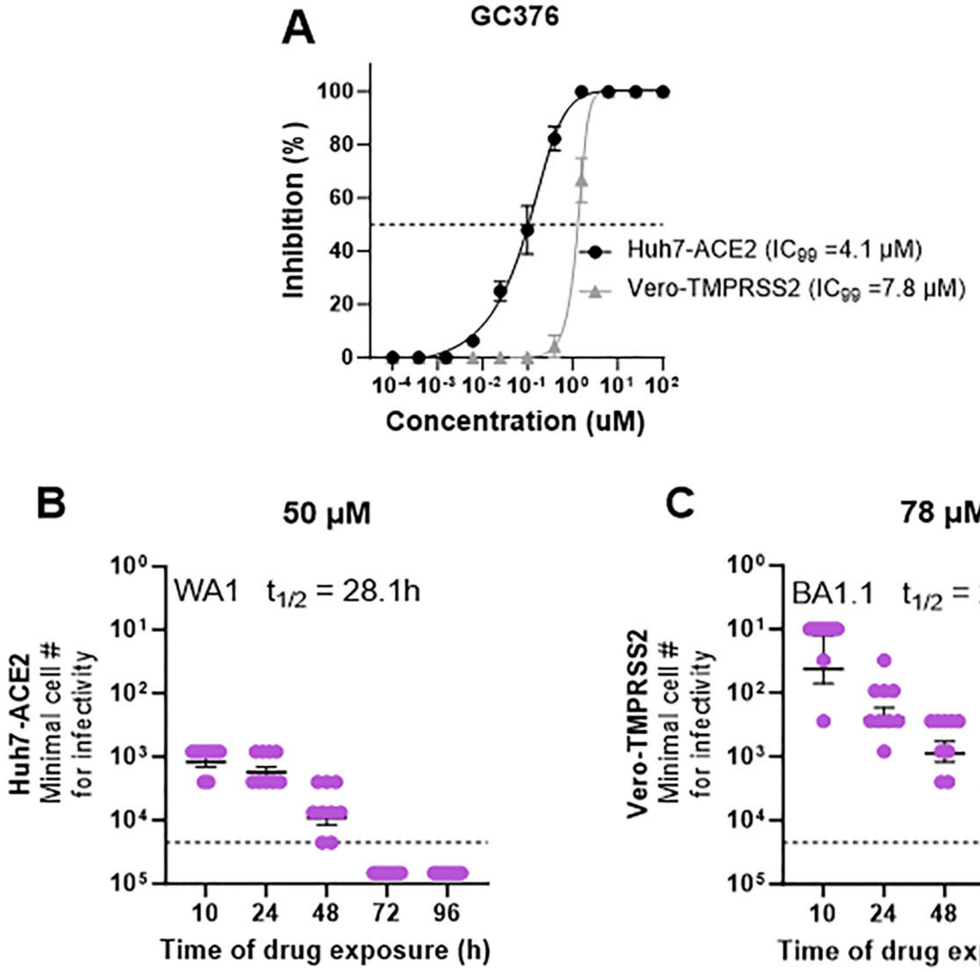


**Figure S2:Testing for SARS-CoV-2 persistence during concurrent treatment with another protease inhibitor (GC-376).** A) Dose-response curves for GC-376 were measured to establish the $IC_{99}$ values of the drug against ic-SARS-CoV-2/WA1-mNG in Huh7-ACE2 cells and against Omicron BA.1.1 in Vero-TMPRSS2 cells. B) Endpoint titers for Huh7-ACE2 cells treated with 50 μM GC-376 (10X $IC_{99}$) at each time point of drug withdrawal. Half-life calculated using linear regression from data points that are quantifiable (≥LOQ of assay) shown in the inset text. C) Endpoint titers for Vero-TMPRSS2 cells treated with 78 μM GC-376 (10X $IC_{99}$) at each time point of drug withdrawal. Half-life calculated using linear regression shown in the inset text. Note: A549-ACE2 cells resulted in no virus suppression at 10-fold dosing once drug is withdrawn likely due to weak inhibitory activity of the drug in the cell with higher $IC_{99}$ values (13.4 μM). Therefore, it is not suitable for endpoint titration. **See also Figure 1.**

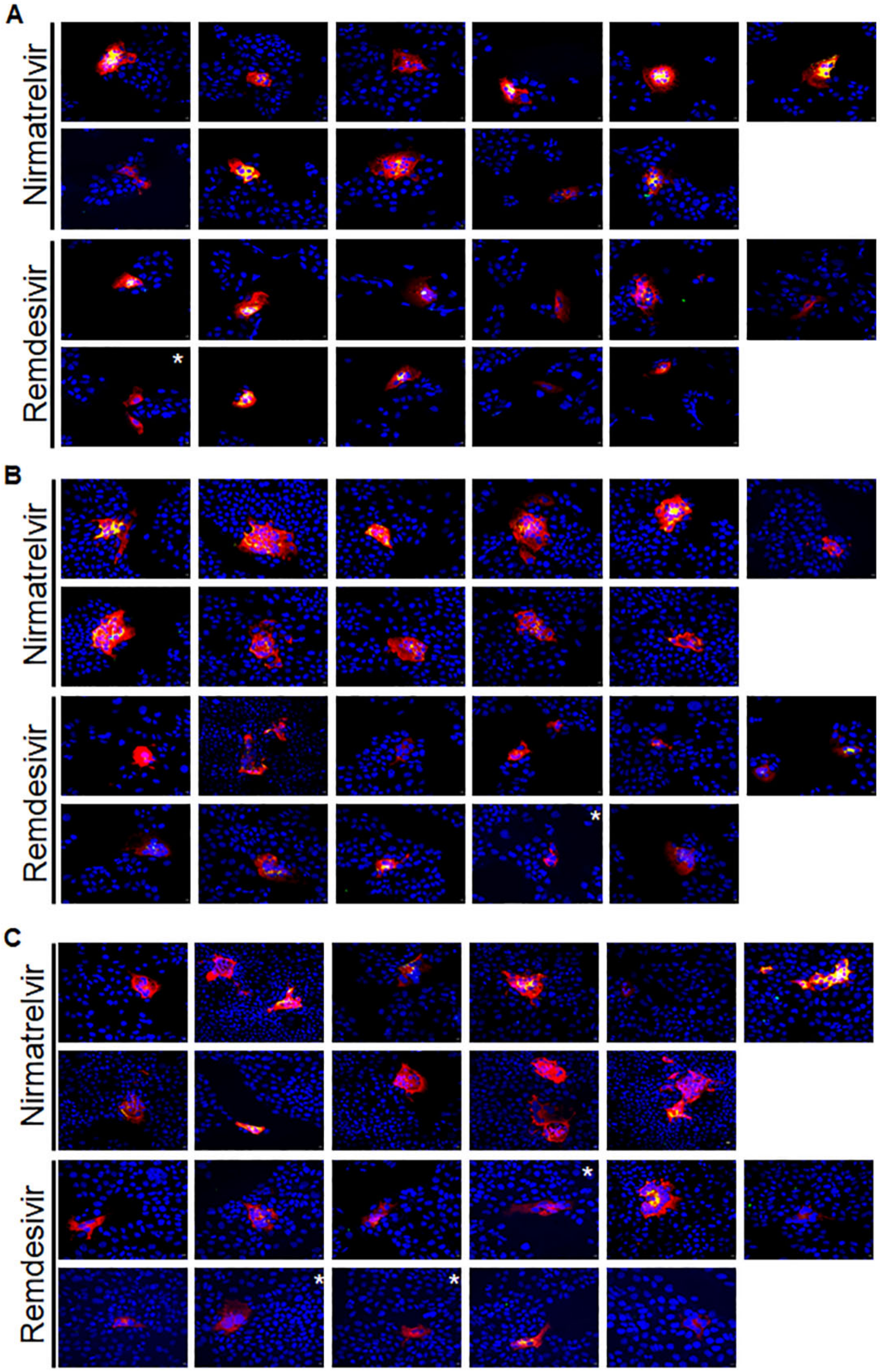


**Figure S3. Simultaneous detection of SARS-CoV-2 genomic RNA and nucleocapsid protein in Huh-7 infected cells post nirmatrelvir or remdesivir treatment.** Huh7-ACE2 were infected with SARS-CoV-2 at 0.5 MOI for 6 hours after which the virus was removed and replaced with growth media supplemented with either 20 μM nirmatrelvir or 1μM remdesivir. At 24 hours (A), 48 hours (B) or 72 hours (C), cells were fixed and processed to detect the genomic viral RNA (green) as well as the viral nucleocapsid protein (red) using RNA-FISH with specific probes together with immunoflurecsence using anti- SARS-CoV-2 nucleocapsid antibodies. Cell nucleus was stained with DAPI (blue). White star indicate higher exposure setting in the red chanel to detect weak nucleocapsid signal. Scale bar = 10μm. **See also Figure 2.**

**KEY RESOURCES TABLE**

| REAGENT or | SOURCE | IDENTIFIER |
|---|---|---|
| **Experimental Models: Cell Lines** | | |
| Huh7-ACE2 | Liu, H. et al (2022) | N/A |
| A549-ACE2 | Ikhlas, S. et al (2022) & BEI Resources | Cat # NR-53821 |
| Vero-TMPRSS2 | BEI Resources | Cat # NR-54970 |
| Vero E6 | ATCC | Cat # CRL-1586<br>RRID:CVCL_0574 |
| **Virus strains** | | |
| ic-SARS-CoV-2/WA1-mNG (ScoV-2/mNG) | Xie, X. et al (2020)<br>World Reference Center for Emerging Viruses & Arboviruses (at UTMB) | N/A |
| BA.1.1 (Omicron) | BEI Resources | Cat # NR-56475 |
| **Reagents** | | |
| Nirmatrelvir/ PF-07321332 | Medkoo Biosciences | Cat # 555985<br>CAS# 2628280-40-8 |
| Ensitrelvir/ S-217622 | Medkoo Biosciences | Cat # 465984<br>CAS# 2647530-73-0 |
| GC-376 | Medkoo Biosciences | Cat # 555227<br>CAS# 1416992-39-6 |
| Remdesivir | Medkoo Biosciences | Cat # 329511<br>CAS # 1809249-37-3 |
| anti- SARS-CoV-1/2 NP antibody | Sigma | Cat# ZMS1075<br>RRID: AB_2893327 |
| Alexa Fluor 647 goat anti-mouse antibody | Thermo Fisher Scientific | Cat # A21236<br>RRID: AB_2535805 |
| **Software** | | |
| GraphPad Prism v9 or v10 | GraphPad Prism Software, Inc.<br>www.graphpad.com | RRID:SCR_002798 |
| Stellaris Probe Designer v 4.2 | LGC BioSearch Technologies<br>https://www.biosearchtech.com/stellaris-designer | N/A |
| SlideBook v 6.0.24 | Intelligent Imaging Innovations<br>https://www.intelligent-imaging.com/slidebook | RRID:SCR_014300 |
| BioRender | https://www.biorender.com/ | RRID:SCR_018361 |
| **Instrumentation** | | |
| Zeiss Axio Observer Z1 microscope | Carl Zeiss Microscopy | N/A |

| | | |
|---|---|---|
| ECHO Revolve fluorescence microscope | ECHO laboratory https://discover-echo.com/revolve/ | N/A |

#### RESOURCE AVAILABILITY

##### Lead Contact

##### Materials Availability

All reagents used in this study are available from the Lead contact and may require completion of a Materials Transfer Agreement prior to sharing.

##### Data and Code Availability

Materials used in this study will be made available but may require execution of a materials transfer agreement. Source data are provided herein.

This paper does not report an original code.

#### EXPERIMENTAL PROCEDURES

##### Cell lines & virus strains

Huh7 (human hepatocellular carcinoma) cells overexpressing human ACE2 cells were prepared by transducing Huh7 cells with lentivirus encoding human ACE2 and selecting for stable expression with 5 µg/mL blasticidin[21] and cultured in Essential Minimal Eagle's medium (EMEM) containing 10% fetal calf serum (FCS) prior to use in dose–response and infectivity assays. Human lung tissue derived cells, line A549 obtained from ATCC (CCL-185) were engineered to overexpress the human angiotensin converting enzyme (ACE2) by stable transfection of a hu-ACE2 expressing lentiviral construct under puromycin selection[22] and cultured in EMEM with 10% FCS and obtained from BEI Resources (NR-53821). *Cercopithecus aethiops* kidney epithelial cells expressing transmembrane protease serine 2 and human ACE2 (Vero-TMPRSS2) were obtained from BEI Resources (NR-54970). An infectious clone of SARS-CoV-2 WA1 containing the mNeonGreen reporter (ic-SARS-CoV-2/WA1-mNG) was obtained from the World Reference Center of Emerging Viruses and Arboviruses (WRCEVA) at the University of Texas Medical Branch[17] and propagated using Vero E6 cells at the high containment laboratory at Aaron Diamond AIDS Center at Columbia University Medical Center. The resultant progeny virions were titered in Vero E6 cells prior to use in dose-response and infectivity experiments. Omicron BA.1.1 isolate hCoV-19/USA/HI-CDC-4359259-001/2021 was procured from BEI Resources (NR-56475), passaged in Vero-TMPRSS2 cells and titered using Reed & Muench method prior to performing infectivity experiments in these cells.

**Drugs**

Nirmatrelvir (PF-07321332), ensitrelvir (S-217622), GC-376, and remdesivir were purchased from Medkoo Biosciences. All drugs were >96% pure and dissolved in DMSO prior to using in dose-response and infectivity experiments.

**Dose-response inhibition of virus**

Triplicates of serial (1 in 4) dilution for each drug starting at 100 µM concentration was made in EMEM (+10% FCS) and plated on the surface of a monolayer of 20,000 target cells per well seeded overnight, at 100 µL volumes in a 96-well plate format. SARS-CoV-2 was added 10 to 15 min later at 0.5 MOI per well. Cells were incubated at 37°C/5%$CO_2$ for 70h prior to measuring cytopathic effects (CPE) under the microscope. CPE was determined relative to the wells lacking any treatment (virus controls) and plotted as a function of the dose using non-linear asymmetric five-parameter dose-response curve in GraphPad Prism version 9.4 to determine the corresponding 50% and 99% inhibitory concentrations of the drug in the cell line. Dose-response curves were performed at least three times in each cell line to determine the IC99 values of all the drugs.

**Determination of minimal cell number for recovering infectivity**

Virus endpoint titrations were performed to determine the efficacy of protease and polymerase inhibitors at doses 10-fold or higher than their 99% inhibitory concentrations in two or more cell lines. Three formats were used to determine the infectivity levels in vitro, referred to as pre-treatment, concurrent treatment, or post-treatment.

For pre-treatment formats, nirmatrelvir, ensitrelvir or remdesivir was added to a monolayer of Huh7-ACE2 cells at doses 10-fold of their $IC_{99}$ for 6 h prior to infecting them with ic-SARS-CoV-2/WA1-mNG at 0.5 MOI. Cells were incubated with drugs and/or virus at $37^oC/5\%CO_2$ for 24 /48/ 72 h. At each time point, media containing drug and/or free virions were removed, and cells were washed 3x with phosphate-buffered saline. Post-washes, the cells were trypsinized with 0.25% Trypsin-EDTA ($37^oC/5\%CO_2$ for 3 min) and collected in EMEM with 10% FCS. Cells were centrifuged and washed again before the cell pellet was re-suspended into 200 -250 µL of EMEM and counted using Bio-Rad TC10 automated cell counter. Counted cells were serially diluted 3-fold in EMEM to set up an endpoint titration in 96-well plates starting at 22,500 cells/well and 100 µl of each dilution was overlaid on Vero cells expressing ACE2 and TMPRSS2 (Vero-TMPRSS2), which are extremely sensitive to virus replication and exhibit prominent CPE. The dilutions were incubated at 37C/5%CO2 for 72h prior to determining the viral endpoint titer from each of the treated concentrations of the extract. Three replicates of titration were set up for each drug for each experiment. Each experiment was performed in triplicates and the minimal cell number from an experiment is plotted for each time point at which drug was removed from the medium.

For the concurrent treatment format of experiments, we used three different cell lines for the study. Huh7-ACE2 and A549-ACE2 human cell lines of liver or lung origin were infected with ic-SARS-CoV-2/WA1-mNG at 0.5 MOI for 10 min prior to adding nirmatrelvir, ensitrelvir, GC376 or remdesivir at concentrations of 10-fold or higher than their $IC_{99}$. A more rigorous time course was used for this study by performing removal of drug at 10, 24, 48, 72 and 96h post infection. The drug removal was performed similarly to the pre-treatment format and three-fold endpoint dilutions starting at 22500 cells/well were placed into a monolayer of Vero-TMPRSS2 cells. For infection of Omicron BA.1.1 isolate, Vero-TMPRSS2 cells were infected at 0.5 MOI for 10 min prior to drug treatment as above. Following drug-removal, the end-point titration was performed using 3-fold dilutions of the collected cells onto a new monolayer of Vero-TMPRSS2 cells. The diluted cells were incubated at 37C/5%CO2 for 72h prior to determining the viral endpoint titer from each of the treated concentrations of the extract. Three replicates of titration were set up for each drug for each experiment. Each experiment was performed in triplicates, and the minimal cell number needed to recover infectious virus from an experiment is plotted for each time point when the drug was removed from the medium.

For the post-infection treatment format, Huh7-ACE2 cells were infected with ic-SARS-CoV-2/WA1-mNG at 0.5 MOI for 6h in 37C/5%CO2. Thereafter, the infection medium was replaced with a medium containing nirmatrelvir or remdesivir at a concentration of 10-fold higher than its 99% inhibitory dose in the cells. Cells were incubated at 37C/5%CO2 for 24, 48 or 72h after which the cells were washed off the drug and free virion, trypsinized, washed again, and counted. The collected cells were then diluted 3-fold starting at 2500 cells/well and overlaid onto a fresh monolayer of Vero-TMPRSS2 cells. Three replicates of each titration were set up for each drug, and each experiment was performed in triplicates. Minimal cell numbers needed to yield infectious virus obtained from the experiment were plotted for each time point when the drug was removed.

**Single molecule RNA FISH and immunofluorescence**

Simultaneous detection of intracellular SARS-CoV-2 viral genomic RNA and nucleocapsid protein using immunofluorescence combined single molecule RNA FISH was performed as described by Kochan at el[20] with minor modifications. Sterile 13mm round glass coverslips were placed in a 12-well plate and 75,000 Huh7-ACE2 cells were plated in 1ml DMEM growth media per well, 24 h prior for infection and drugs treatment. At every time point, the cells were washed 3 times with 1x PBS and fixed using 4% formaldehyde in PBS (10 min) followed by additional 3 washes with 3 times with 1x PBS. Samples that were taken at the 24 and 48 h time points were stored at 4$^{o}$C and processed together with the 72 h samples. For nucleocapsid detection at the immunofluorescence step, mouse monoclonal anti- SARS-CoV-1/2 NP Antibody (clone 1C7C7; Sigma Cat# ZMS1075) at a concentration of 1µg/ml was used as the primary antibody, and Alexa Fluor 647 goat anti-mouse (ThermoFisher Scientific Cat# A21236) at a concentration of 2µg/ml was used as the secondary antibody. For single-molecule RNA FISH to detect the positive strand of the SARS-CoV-2 genome, probe sequences were designed using Stellaris Probe Designer version 4.2 (https://www.biosearchtech.com/stellaris-designer) with the following parameters: organism, human; masking level, 5; oligo length, 20 nt; minimum spacing length, 3 nt and the nucleotide sequence of ORF1a from the ic-SARS-CoV-2/WA1-mNG SARS-CoV-2[17] similarly to previous published study[23]. Oligonucleotide probes were purchased labeled with TAMRA dye and used at a 250nM final concentration.

**Microscopy**

Images were acquired using motorized spinning-disc confocal microscope (Yokogawa CSU-X1 A1 confocal head and Zeiss Axio Observer Z1 microscope) using x63 oil (1.4 NA) and x20 dry (0.8 NA) objectives. Three-dimensional (3D) z stacks were acquired for cells displaying nucleocapsid signal and images were analyzed with SlideBook software (Intelligent Imaging

Innovations; SlideBook version 6.0.24 (40297)). For figures, two-dimensional (2D) projection images of the maximum pixel intensity over the z axis were generated.

**Viral genomic RNA and nucleocapsid quantification**

Intensity measurement of viral nucleocapsid or viral gRNA were achieved by intensity-based segmentation using manually defined thresholding in SlideBook Software. Signal intensity distribution was plotted as a violin plot and statistical significance was determined by one-tailed Student's t test GraphPad Prism version 9.4.

#### QUANTITATION AND STATISTICAL ANALYSIS

**Linear regression analysis to measure decay half-life of infectious virus**

The minimal cell numbers from all the replicates that are required to infect Vero-TMPRSS2 indicator cells at each time point were used to generate a linear regression curve for each drug in each experiment. The reciprocal of the cell number for each of the 9 replicates for every drug was averaged to determine the average frequency of infectivity per time point per drug. The slope of the decay of the average frequency was calculated using a simple linear regression analysis in GraphPad Prism version 10.0 to determine the half-life ($t_{1/2}$) of the infectious material when treated with that drug at a particular dose. A comparison of the half-life of each drug at every dose was calculated and tabulated. Only those points that were quantifiable (≥LOQ of assay) were used to calculate the slope and the corresponding half-lives were represented as less than the obtained number.

**Statistical analysis for measuring significance between groups**

Statistics for significance between test groups in each experiment when shown were performed using the one-tailed student's t-test in GraphPad Prism.

#### References (Supplementary methods)


21. Liu, H., Iketani, S., Zask, A., Khanizeman, N., Bednarova, E., Forouhar, F., Fowler, B., Hong, S.J., Mohri, H., Nair, M.S., et al. (2022). Development of optimized drug-like small molecule inhibitors of the SARS-CoV-2 3CL protease for treatment of COVID-19. Nat Commun *13*, 1891. 10.1038/s41467-022-29413-2.

22. Ikhlas, S., Usman, A., Kim, D., and Cai, D. (2022). Exosomes/microvesicles target SARS-CoV-2 via innate and RNA-induced immunity with PIWI-piRNA system. Life Sci Alliance *5*. 10.26508/lsa.202101240.

23. Lee, J.Y., Wing, P.A.C., Gala, D.S., Noerenberg, M., Jarvelin, A.I., Titlow, J., Zhuang, X., Palmalux, N., Iselin, L., Thompson, M.K., et al. (2022). Absolute quantitation of individual SARS-CoV-2 RNA molecules provides a new paradigm for infection dynamics and variant differences. Elife *11*. 10.7554/eLife.74153.
